# Sufficiency analysis of estrogen responsive enhancers using synthetic activators

**DOI:** 10.1101/479709

**Authors:** Matthew Ginley-Hidinger, Julia B. Carleton, Adriana C. Rodriguez, Kristofer C. Berrett, Jason Gertz

## Abstract

Multiple regulatory regions bound by the same transcription factor have been shown to simultaneously control a single gene’s expression. However, it remains unclear how these regulatory regions combine to regulate transcription. Here we test the sufficiency of promoter-distal estrogen receptor α (ER)-binding sites (ERBS) for activating gene expression by recruiting synthetic activators in the absence of estrogens. Targeting either dCas9-VP16(10x) or dCas9-p300(core) to ERBS induces H3K27ac and activates nearby expression in a manner similar to an estrogen induction, with dCas9-VP16(10x) acting as a stronger activator. The sufficiency of individual ERBS is highly correlated with their necessity, indicating an inherent activation potential that is associated with the binding of RNA polymerase II and several transcription factors. By targeting ERBS combinations, we found that ERBS work independently to control gene expression when bound by synthetic activators. The sufficiency results contrast necessity assays that show synergy between these ERBS, suggesting that synergy occurs between ERBS in terms of activator recruitment, whereas directly recruiting activators leads to independent effects on gene expression.

## Introduction

Gene expression enhancers are genomic loci that act in *cis* to control a distal gene’s expression level. There are two orders of magnitude more predicted enhancers in the human genome compared to gene promoters (Roadmap Epigenomics Consortium et al., 2015), indicating that many mammalian genes are regulated by multiple enhancers. Analysis of 3D genome architecture (ENCODE Project Consortium, 2012) and the expression of enhancer RNAs (Andersson et al., 2014) corroborate the idea that the average human gene is regulated by the combined action of many enhancers. Functional studies into enhancer combinations have found that enhancers can work together in an additive/independent (Bender et al., 2012; Fujioka et al., 1999), synergistic (Lam et al., 2015; Torbey et al., 2018), or redundant (Hong et al., 2008; Osterwalder et al., 2018) manner, indicating that enhancers can combine to regulate gene expression in complex and diverse ways.

There are two different approaches to functionally perturb enhancers to study enhancer function: necessity and sufficiency. Deleting or inhibiting enhancer function tests the necessity of an enhancer for endogenous gene expression. The sufficiency of enhancer sequences can be studied by ectopic reporter assays (Catarino and Stark, 2018). However, testing only enhancer sequences does not uncover how enhancers act in their endogenous environment. To determine whether an enhancer region is sufficient within the genomic context, enhancers must be directly activated in an unbiased way. Most studies of enhancer function and combinatorics involve genetic deletion of the region(s) of interest. Genetic deletion, along with CRISPR interference-based approaches (Carleton et al., 2017; Fulco et al., 2016; Gilbert et al., 2013; Korkmaz et al., 2016; Thakore et al., 2015), test the necessity of a genomic region or combination of genomic regions for regulatory activity. Regulatory element screens that test sufficiency identify shared and unique regulatory elements in comparison to screens for necessity, indicating that necessity and sufficiency are complementary assays (Klann et al., 2017). Testing sufficiency of an enhancer by direct activation provides additional insight into the innate capabilities of an enhancer, the requirements of enhancer function and whether genomic properties at enhancer regions associate with an enhancer’s potential for modulating transcription. However, testing the sufficiency of genomic regulatory regions in their native context is less commonly undertaken than necessity.

Reporter assays, in which a plasmid containing a potential enhancer sequence controls the expression of a reporter gene, are one way of testing enhancer sufficiency (Catarino and Stark, 2018). However, this technique does not analyze an enhancer within its native chromatin context and is likely missing information that might be critical for the ability of an enhancer to regulate gene expression, such as epigenetic markers, interaction of enhancers with specific promoters and properties of adjacent genomic regions (Cunningham et al., 2018). CRISPR activation (CRISPRa) is a recently developed tool for activating gene expression from specific regions of the genome. CRISPRa involves the fusion of a catalytically dead Cas9 protein (dCas9) to an activation domain. Different activation domains can be fused to dCas9, including the transcriptional activation domain of VP16 (Cheng et al., 2013) and the core domain of p300 (Hilton et al., 2015). VP16 is a herpes simplex virus transcription factor which recruits a variety of host factors, including general transcription factors, mediator, and histone acetyltransferases (Hirai et al., 2010). p300 is a histone acetyltransferase linked with activation of many genomic regions (Delvecchio et al., 2013). CRISPRa provides a strategy for turning on specific genes when targeted to promoter regions with guide RNAs (Chavez et al., 2015; Cheng et al., 2013; Perez-Pinera et al., 2013). CRISPRa can also be targeted to distal regulatory regions in order to test their sufficiency in promoting gene expression (Hilton et al., 2015; Klann et al., 2017; Thormann et al., 2018); however, very few distal regulatory regions have been interrogated in this manner and it is unclear how targeting CRISPRa to combinations of enhancers will impact gene expression.

Many enhancers are controlled by the activity of inducible transcription factors. Estrogen receptor ⍰ (ER) is a steroid hormone receptor which only has gene regulatory activity when it is bound by estrogens. ER acts mostly as an activating transcription factor, binding to thousands of genomic loci and regulating hundreds of genes (Gertz et al., 2012, 2013). The majority of genes that are up-regulated by estrogen have multiple ER bound sites nearby and we previously found evidence of collaboration between ER bound sites in regulating the gene expression response to estrogens (Carleton et al., 2017). Using a CRISPRi-based approach, enhancer interference (Enhancer-i), we found synergistic and hierarchical relationships involving ER bound sites. These relationships were discovered by measuring the necessity of each ER bound site individually and in combination. It is unclear whether ERBS collaboration is a property of the ER transcription factor or dependent on the genomic properties of ER binding sites (ERBS), necessitating an investigation of ERBS sufficiency in the genomic context.

Here, we use CRISPRa to target combinations of regulatory regions that are normally bound by ER (Figure 1A). By targeting these regions in the absence of estrogens, we sought to determine if CRISPRa synthetic activators could recreate the transcriptional response to estrogens. We find that dCas9-VP16(10x) fusion can recreate most of the estrogen response at the four genes tested, while dCas9-p300(core) was not as effective. Targeting CRISPRa to individual regulatory regions and combinations of loci uncovered an additive/independent relationship between sites, in contrast to our previous necessity findings. Our results indicate that ER binding to neighboring enhancers works in a synergistic fashion, but synthetic activators directly recruited to loci normally bound by ER work independently to regulate gene expression.

**Figure 1.**
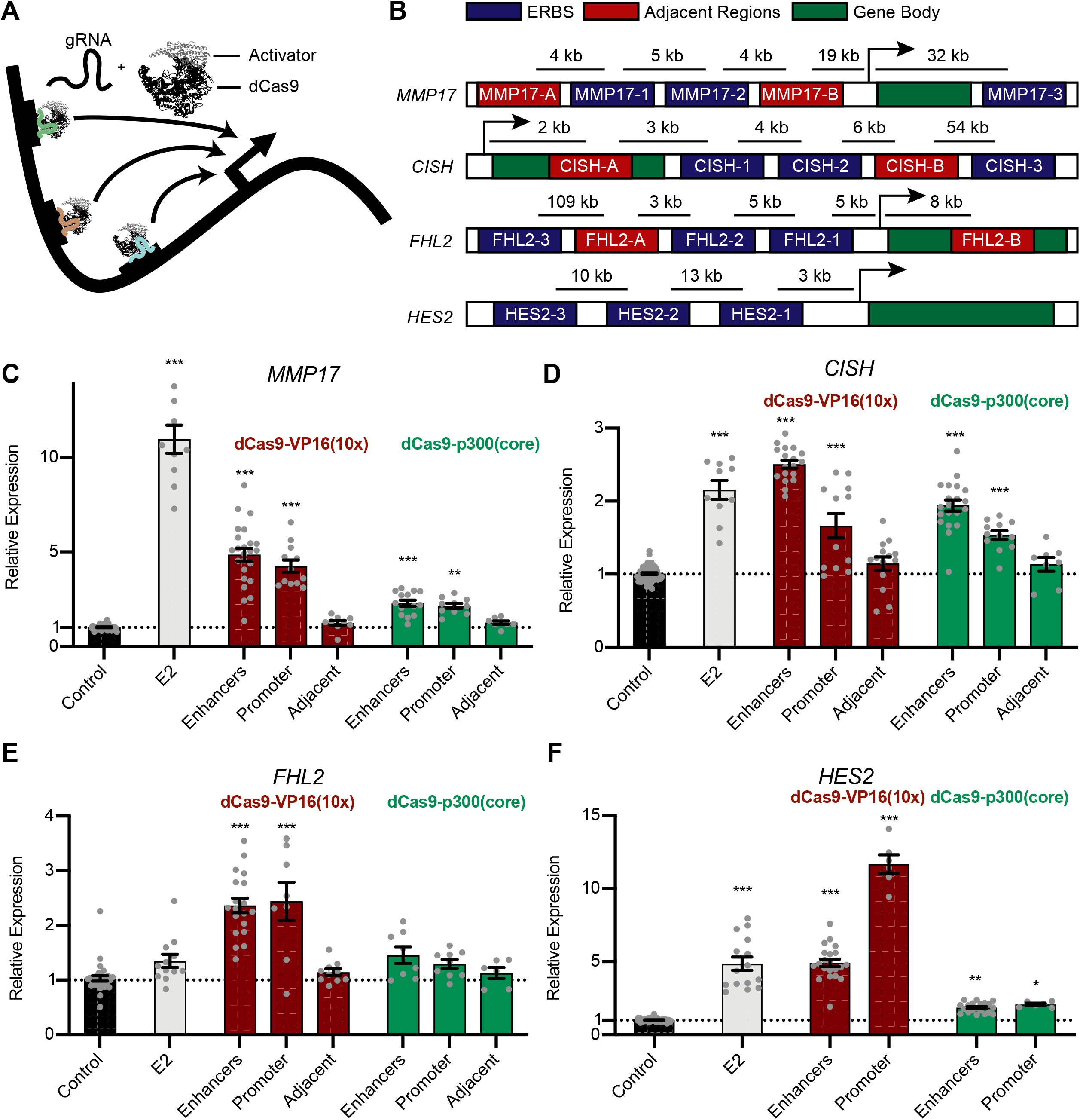
Targeting multiple ERBS with synthetic activators can activate gene expression. **(A)** Cartoon depicting the targeting of multiple ERBS in combination to study combinatorial effects on gene expression. **(B)** Relative locations of ERBS (blue), ERBS-adjacent regions (red) and genes (green) tested in this study. **(C-F)** Targeting all 12 ERBS in combination with dCas9-vp16(10x) (red) or dCas9-p300(core) (green) activated gene expression at *MMP17* **(C)**, *CISH* **(D)**, *FHL2* **(E)** and *HES2* **(F)** to levels that are comparable to an 8-hour E2 treatment (light gray). Targeting all ERBS had significantly higher activation than targeting ERBS-adjacent regions, which is not significantly different than controls that target the *IL1RN* promoter. Error bars represent SEM. P-values are calculated with respect to control using a one-way ANOVA with Dunnett’s multiple comparisons (*: p-value < 0.05, **: p-value < 0.01, ***: p-value < 0.001).

## Results

### Targeting CRISPRa to loci normally bound by ER can mimic transcriptional responses to estrogens

In order to determine if synthetic activators are sufficient to drive an estrogen-like transcriptional response, we evaluated two CRISPRa fusion proteins: dCas9-VP16(10x) (Cheng et al., 2013), which is commonly used at promoters, and dCas9-p300(core) (Hilton et al., 2015), which can activate gene expression from enhancers. Each fusion was expressed from identical expression vectors (see Methods, Figure S1A) in order to directly compare their ability to activate gene expression from distal regulatory regions normally bound by ER. We first targeted dCas9-VP16(10x) and dCas9-p300(core) to the *IL1RN* promoter, a gene which can be highly activated by CRISPRa (Cheng et al., 2013; Hilton et al., 2015; Perez-Pinera et al., 2013), and observed a similar level of activation for *IL1RN* with both dCas9 fusion constructs in Ishikawa cells, an endometrial cancer cell line (Figure S1B).

The dCas9 fusions were targeted to a pool of ER binding sites (ERBS) to determine if the estrogen response could be recapitulated with CRISPRa. We chose a set of 12 ERBS that were within 100 kilobase pairs (kb) of 4 genes that are normally responsive to estrogen. *MMP17* has two upstream and one downstream ERBS, *CISH* has three downstream ERBS, and *FHL2* and *HES2* both have three upstream ERBS (Figure 1B). We targeted the CRISPRa fusions to the 12 ERBS simultaneously and measured gene expression of the target genes in the absence of estrogens and therefore the absence of ER binding. We observed significant gene expression activation by dCas9-VP16(10x) at all 4 genes tested, while activation by dCas9-p300(core) was significant at 3 genes (*MMP17*, *CISH* and *HES2*) (Figure 1C-F). For comparison, we also targeted our constructs to the promoter regions of these 4 genes and found that targeting ERBS often results in equal or greater activation than targeting the promoter (Figure 1C-F). dCas9-VP16(10x) was consistently a stronger activator than dCas9-p300(core). The level of gene expression driven by dCas9-VP16(10x) was somewhat correlated with the fold change in gene expression seen with a 17β-estradiol (E2) induction at this set of four genes (r = 0.8) (Figure S1C). These results demonstrate that CRISPRa targeted to ERBS can mimic the activation seen in an E2 transcriptional response.

To determine if the activation potential is specific to ERBS, we targeted dCas9-VP16(10x) and dCas9-p300(core) to a total of 6 regions surrounding *MMP17*, *CISH* and *FHL2* that are at most 8 kb away from ERBS discussed above (or the TSS in the case of FHL2-B) (Figure 1B). Regions with low DNase I hypersensitivity signal (Gertz et al., 2013) were chosen, in order to limit the probability that the locus was an active regulatory region controlled by other transcription factors. As *HES2* is in a highly active region with several DNase I hypersensitive sites and histone H3 lysine 27 acetylation positive loci, multiple nearby genes and many transcription factor binding events, we were unable to choose sites that were not potential regulatory regions at this locus. In choosing adjacent regions, we aimed to keep the distance to the TSS similar without being too close to the ER bound site. We observed some transcription factors binding to the chosen adjacent regions, notably at CISH-A (Table S3). When targeting the ERBS adjacent sites, we did not observe significant activation over the control of targeting the *IL1RN* promoter (Figure 1C-E). The inability of ERBS adjacent regions to regulate gene expression with synthetic activators indicates specificity when testing sufficiency of regulatory regions with CRISPRa and differences in activation potential between ERBS and nearby non-ERBS.

### dCas9 activator fusions can target precise genomic loci and induce histone acetylation

To test whether CRISPRa is successfully targeted to the intended genomic regions surrounding our genes of interest, we conducted a Chromatin Immunoprecipitation followed by sequencing (ChIP-seq) experiment using an antibody that recognizes an HA epitope tag on dCas9. dCas9-VP16(10x) was targeted to 19 loci, consisting of 12 ERBS, 6 ERBS-adjacent regions and the *IL1RN* promoter (Figure 1B). At all of these loci, we observed a distinct HA (dCas9) signal at the targeted site when compared to non-targeted controls (Figures 2A,B and S2A-C). Additionally, dCas9-p300(core) was successfully targeted to all 12 ERBS (Figure 2A). Successful targeting of dCas9-VP16(10x) to ERBS-adjacent regions indicates the lack of activation from these regions is based on genomic properties of the adjacent regions and not a result of the targeting efficiency.

**Figure 2.**
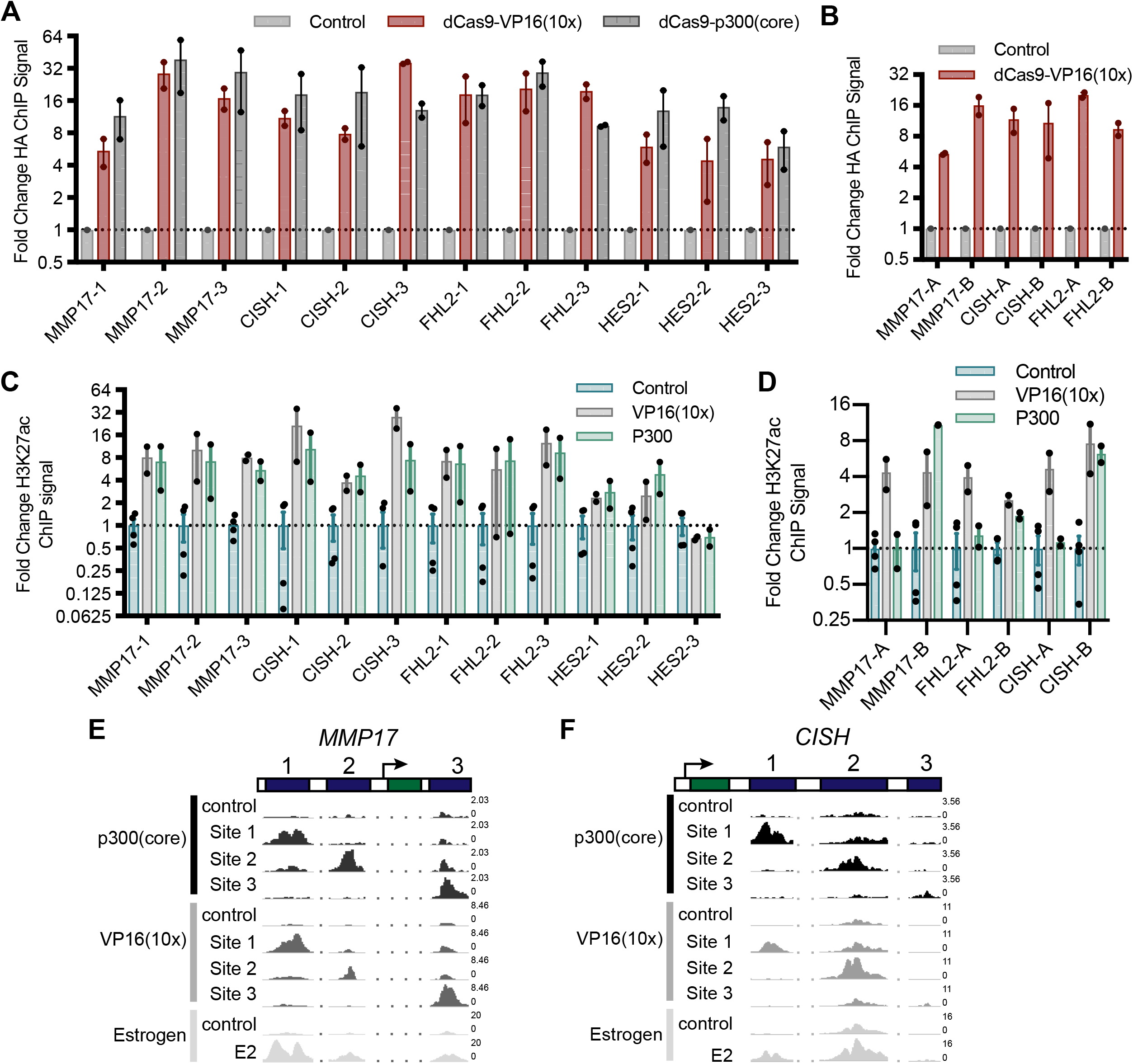
Targeting dCas9-activator constructs is specific and induces H3K27ac. **(A-B)** The relative HA ChIP-seq signal, an epitope tag on dCas9, is shown for all targeted ERBS **(A)** and adjacent regions **(B)** and compared to non-targeted controls. Points indicate individual replicates and error bars represent SEM. **(C)** The fold change induction of H3K27ac ChIP-seq signal across all targeted loci shows no significant difference between dCas9-p300(core) and VP16(10x). **(D)** H3K27ac was induced at all adjacent regions by dCas9-VP16(10x) and at 3 of 6 adjacent regions by dCas9-p300(core). Points indicate individual replicates and error bars represent SEM. **(E-F)** Browser tracks of H3K27ac induced by targeting ERBS with dCas9-p300(core), dCas9-VP16(10x) or an 8-hour E2 treatment are shown at *MMP17* **(E)** and *CISH* **(F)**. Numbers on the right of the tracks indicate the track height in reads per million.

Histone 3 lysine 27 acetylation (H3K27ac) is a histone modification found at active regulatory regions and is directly deposited by p300 (Raisner et al., 2018). We therefore performed ChIP-seq with an antibody that recognizes H3K27ac to determine if the CRISPRa fusions were able to cause H3K27ac at targeted sites. For 18 of 19 targeted loci, we observed increased H3K27ac (Figures 2C,D, S2C-G). Notably, at HES2-1 and HES2-3, there is significant baseline H3K27ac present, possibly due to the binding of other transcription factors to these sites (Table S2). For ERBS, the patterns of acetylation are similar to E2 induced H3K27ac (Figures 2E,F and S2D,E). We observed similar fold changes in H3K27ac when using dCas9-VP16(10x) and when using dCas9-p300(core) targeted to ERBS (Figures 2C,E,F and S2D,E; p-value = 0.108, paired t-test). This result is in contrast to the greater gene activation induced by dCas9-VP16(10x), indicating that histone acetylation of a distal regulatory element is not fully predictive of target gene activation. Consistent with the idea that H3K27ac by itself is not sufficient to drive maximal expression, we observed H3K27ac at the adjacent regions even though they were unable to induce gene expression (Figure S2F). Additionally, we conducted ChIP-seq with an antibody for RNAPII and found limited RNAPII recruitment to ERBS by dCas9-p300(core) or dCas9-VP16(10x) (Figure S2H), though we saw RNAPII recruitment by dCas9-VP16(10x) at the *IL1RN* promoter (Figure S2C).

### dCas9-VP16(10x) activates gene expression from individual ER binding sites

Since targeting dCas9-VP16(10x) to all ERBS simultaneously resulted in gene activation of all genes, we next sought to determine if targeting individual ERBS with dCas9-VP16(10x) is sufficient to increase gene expression. At the *MMP17* locus, all three ERBS activated gene expression above the control level when targeted individually (Figure 3A, pairwise p-values in Table S1). Targeting MMP17-1 resulted in the highest induced expression, MMP17-2 exhibited the weakest activation and MMP17-3 led to an intermediate change in *MMP17* expression. When targeting the ERBS surrounding *CISH*, CISH-1 and CISH-2 induced a similar level of activation, while CISH-3 did not result in activation (Figure 3B). The *FHL2* gene was induced strongest by FHL2-1. *FHL2* was also activated by FHL2-2 and FHL2-3, but to a lower level (Figure 3C). At *HES2*, we observed a high level of activation from HES2-1 and slight activation from HES2-2 and HES2-3 (Figure 3D). Targeting dCas9-p300(core) to individual ERBS resulted in a lower, but correlated, activation than dCas9-vp16(10x) (r = 0.633, Figure S4A-D), indicating that the relative strength of enhancers may be independent of the synthetic activator used, while absolute strength can be controlled by the strength of the synthetic activator.

**Figure 3.**
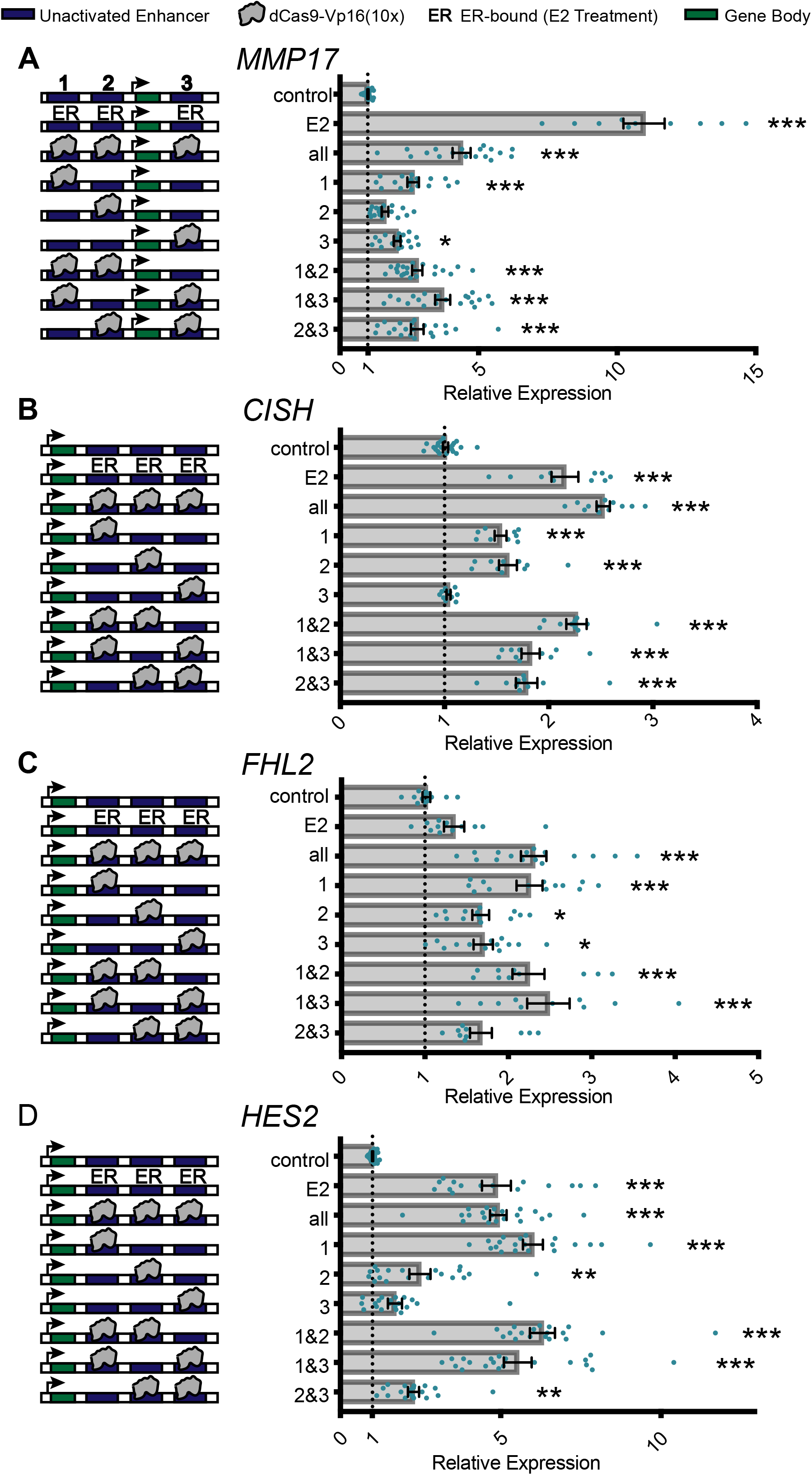
Activation of gene expression by targeting CRISPRa to combinations of enhancers. **(A-D)** (left) The combination of ERBS targeted by dCas9-VP16(10x) or bound by ER upon E2 treatment are shown in a schematic. **(A-D)** (right) The relative fold change in expression as measured by qPCR, when compared to control cells with guides targeting the *IL1RN* promoter, was determined for each combination of ERBS. Each data point is shown as a blue dot, error bars represent SEM. Pairwise log2 ratios and significance levels are given in Table S1 (Pairwise t-test p-values comparing to control: *** < 0.001, ** < 0.01, * < 0.05).

To ensure that the activation observed from individual sites is specific to the targeted site, we tried to activate expression in previously derived Ishikawa lines with homozygous deletion of the targeted ERBS for the *MMP17* and *CISH* sites (Carleton et al., 2017). In each case, no detectable gene activation was observed in the deletion lines indicating specificity in targeting individual ERBS (Figure S3A,B). We theorized the observed differences in activation at individual sites may be due to biases in dCas9 targeting, preferential H3K27ac or differences in RNAPII recruitment. However, we found no significant correlation between expression activation and fold change HA ChIP-seq signal, fold change H3K27ac ChIP-seq signal or fold change RNAPII ChIP-seq signal (Figure S3C-E). The ability to activate gene expression from several individual enhancers adds to a growing body of literature showing that a gene’s expression can be controlled by multiple regulatory regions. Furthermore, the unique level of activation that results from targeting individual enhancers suggests that enhancers do not contribute equally to gene activation, even when bound by a synthetic activator.

### Enhancers bound by synthetic activators work independently to regulate transcription

Based on the observation that multiple ERBS nearby each gene were capable of activating gene expression when targeted individually by dCas9-VP16(10x), we investigated how synthetic activator bound enhancers collaborate to control gene expression by targeting pairs of ERBS. In general, ERBS appeared to combine independently when simultaneously targeted. For example, CISH-1 and CISH-2 are the two strongest individual activators of *CISH* and each increase gene expression to approximately 40% of maximum observed activation while the combination of CISH-1 and CISH-2 increases gene expression to 80% (Figure 3B). A similar pattern was observed for *MMP17* (Figure 3A), while *FHL2* (Figure 3C) and *HES2* (Figure 3D) exhibit sub-additive effects that may represent saturation.

In order to quantitatively understand the interactions between ERBS when bound by a synthetic activator, we created a thermodynamic model of RNA Polymerase II (RNAPII) recruitment (as a surrogate for transcription) by ERBS (Buchler et al., 2003; Shea and Ackers, 1985). To describe the differences in gene expression seen from our combinatorial activation studies, we fit relative energy parameters to an abstracted model of combinatorial synthetic activation (Figure 4A). The model included 4 sets of interactions: (1) Interactions between dCas9-VP16(10x) and targeted ERBS (Figure 4, red), (2) interactions between ERBS-bound synthetic activators (Figure 4, green), (3) interactions between ERBS-bound synthetic activators and RNAPII (Figure 4, blue), and (4) interactions between RNAPII and a gene’s promoter (Figure 4, purple). This model assumes that the probability of RNAPII binding is proportional to a gene’s expression. We used the correlation between the probability of RNAPII being bound and gene expression data from targeting combinations of ERBS (Figure 3) to fit model parameters. We ran the parameter optimization with many random starts, then selected parameters that fit gene expression levels reasonably well (within 0.1 of optimal) (see Methods). We consequently observed a range of parameters that were locally optimal (Figure 4B-E). In some cases, we observed multimodal parameter distributions, which is likely due to parameters balancing each other in different ways, leading to multiple local optima.

**Figure 4.**
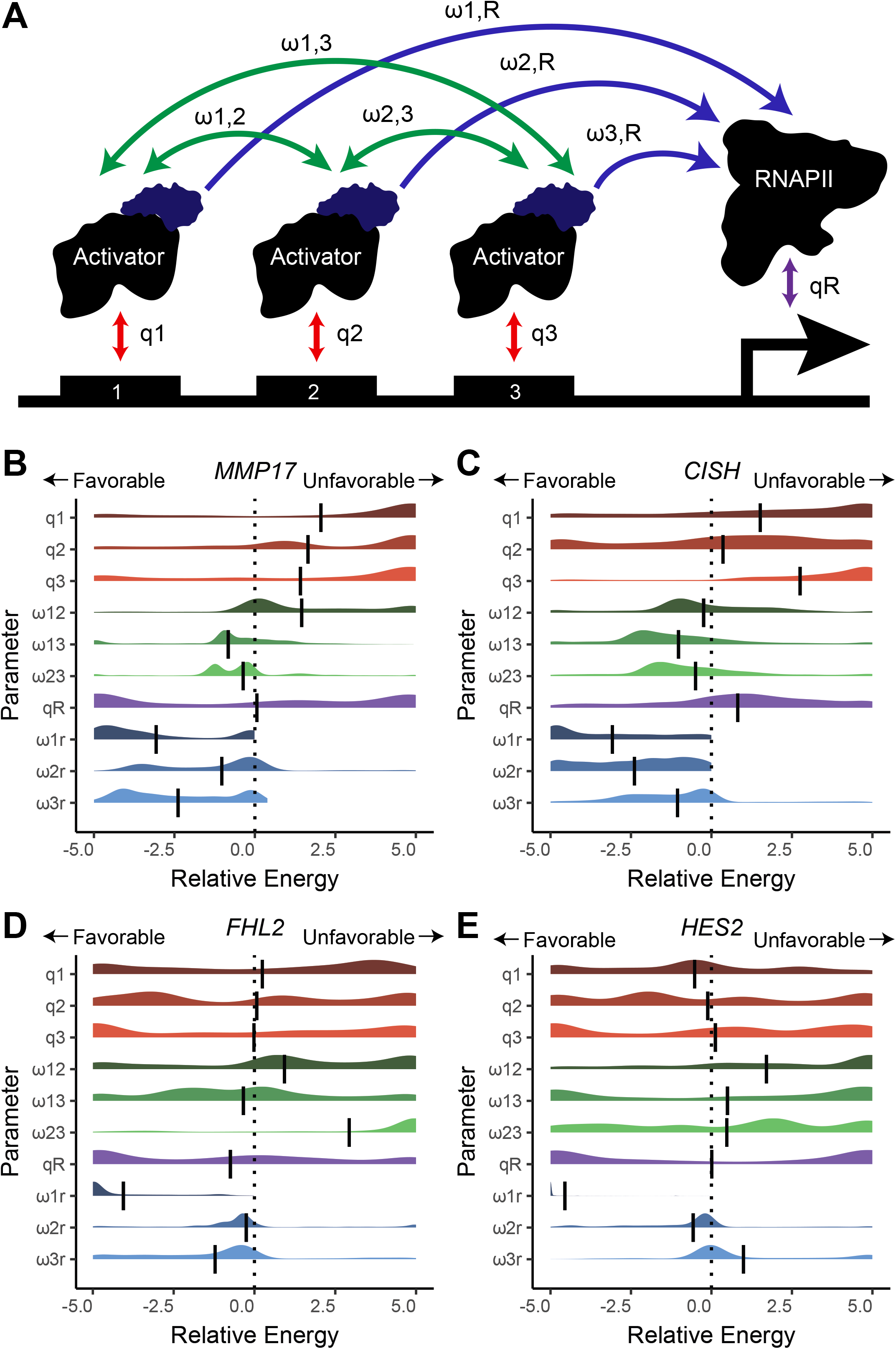
Thermodynamic modelling reveals little cooperativity between synthetic activator bound ERBS. **(A)** Schematic showing the set of modelled parameters. **(B-E)** Parameters were fit to gene expression data for *MMP17*, *CISH*, *FHL2* and *HES2* (from Figure 3). Plots show the distribution of fitted parameters. Parameter sets were selected if the modelled data correlated with gene expression data within 0.1 of an optimal correlation. Vertical bars represent the mean.

In this model, the activation observed from targeting individual loci is largely captured by the interaction terms between RNAPII and synthetic activators (parameter ranges shown in blue in Figure 4), where more favorable (more negative) interactions are indicative of more gene activation. For example, at *FHL2*, site FHL2-1 has the largest impact on expression and the most favorable interaction with RNAPII, while FHL2-2 and FHL2-3 have modest effects on expression and slightly favorable RNAPII-synthetic activator interactions (Figure 4D). In these models, the interaction between synthetic activators and RNAPII can be balanced by differential recruitment of the synthetic activators to the ERBS (shown in red in Figure 4). For example, CISH-1 and CISH-2 both activate gene expression to similar levels, but CISH-1 is modeled with the synthetic activator more strongly recruiting RNAPII while CISH-2 is modeled as binding the synthetic activator with more efficiency (Figure 4C). The relationship between ERBS in the sufficiency experiments is best captured by the interaction terms between synthetic activators bound to ERBS (parameter ranges shown in green in Figure 4). For all studied genes, we do not see strong cooperativity between ERBS. For *MMP17* and *CISH*, we observed relatively neutral interactions between ERBS (Figure 4B,C). The best fits for *FHL2* and *HES2* were mostly competitive models where certain ERBS inhibit others (Figure 4D,E). This may result from a limit in how much gene activation can be driven by the synthetic activators. For example, dCas9-VP16(10x) at HES2-1 results in a similar gene expression level as targeting all ERBS surrounding *HES2* simultaneously. Even though targeting of HES2-2 or HES2-3 has some activity in isolation, they are unable to increase expression beyond HES2-1 targeting (Figure 3D). This is captured in the model as unfavorable interactions between HES2-1 and the other ERBS. In general, the lack of cooperativity in these models supports the conclusion that these sites work independently to activate gene expression when targeted with dCas9-VP16(10x).

It is possible that the observed independence of synthetically activated ERBS is due to the strength of dCas9-VP16(10x), which may override the subtleties of enhancer synergy. We therefore used thermodynamic models to analyze activation by a weaker activator, dCas9-p300(core), at the 3 genes which can be activated by this construct. Again, we observed an independent relationship at *MMP17* (Figure S4E). At *CISH* and *HES2*, we observed bimodal distributions of some interaction parameters, where a subset of fitted models indicate synergy between enhancers; however, the majority of model fits indicate independence between enhancers when bound by dCas9-p300(core) (Figure S4F,G). The presence of multiple local optima might be due to low levels of activation and lower signal-to-noise ratios. Overall, we see that sites activated by dCas9-p300(core) do not display strong patterns of cooperativity, consistent with the results from dCas9-VP16(10x) targeting.

### Comparison between necessity and sufficiency of ERBS

We previously assessed the necessity of the nine ERBS nearby *CISH*, *MMP17*, and *FHL2* in producing a transcriptional response to estrogen using Enhancer-i (Carleton et al., 2017). At these three genes, we found the predominant ERBS for activating gene expression when testing sufficiency was the same as when testing necessity (e.g. MMP17-1 and FHL2-1). In order to normalize the relative impact of targeting individual ERBS for each gene, we calculated the z-score of relative expression when targeting an individual ERBS compared to targeting the other individual ERBS for that gene. We observed a strong correlation between the relative necessity, as measured by Enhancer-i (and validated by genetic deletion), and sufficiency, as measured by dCas9-VP16(10x) targeting (r = 0.840, Figure 5A). The consistent importance of individual ERBS, in terms of sufficiency and necessity, suggests that each ERBS has a native activation potential that is unique to the site.

**Figure 5.**
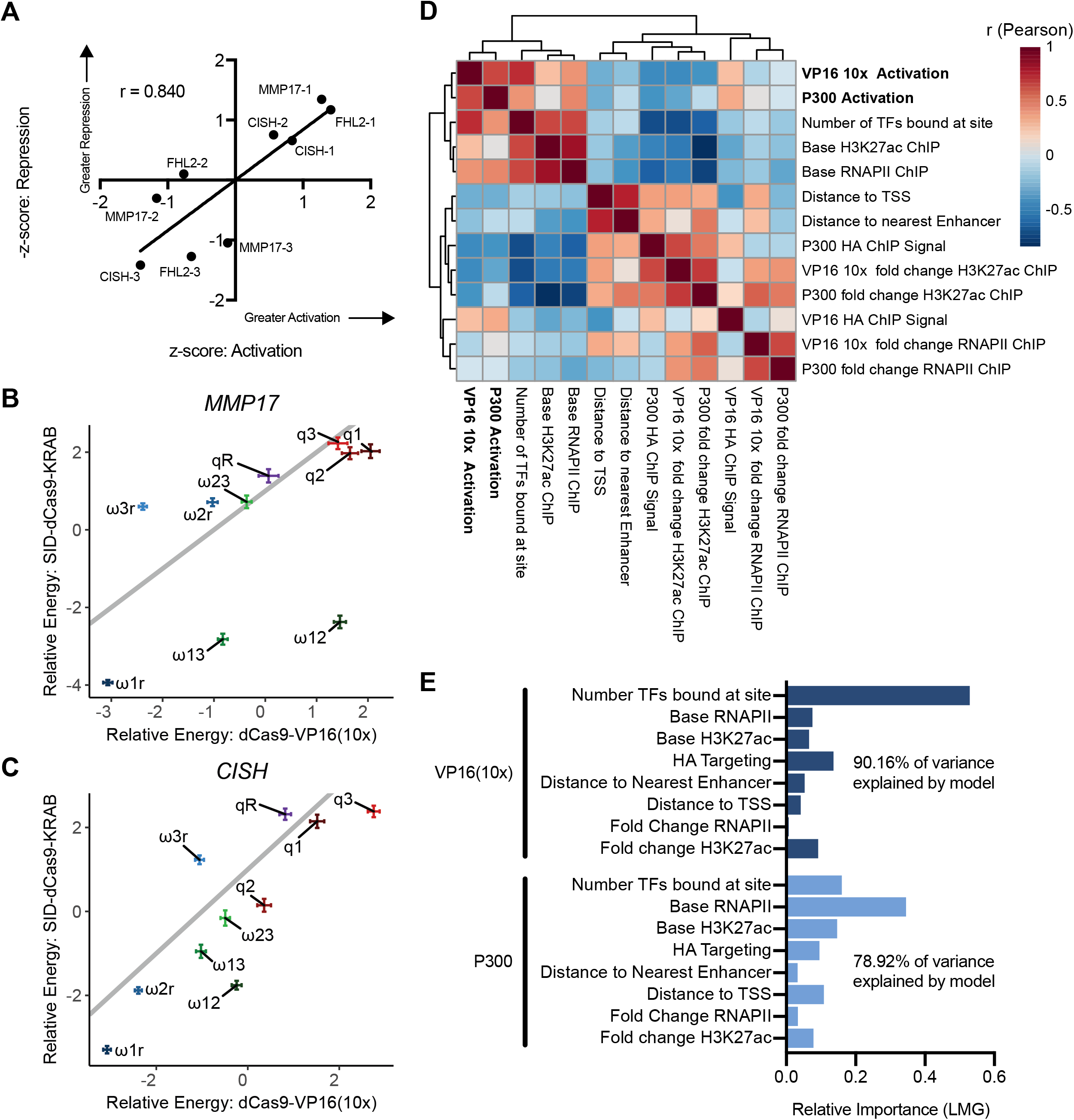
Sufficiency-necessity comparison shows similar results at individual sites, but differences in cooperation between sites. (A) Scatter plot shows relative expression, as measured by z-score, for both activation and interference at individual ERBS. Z-scores were negated for interference so that a higher score is associated with greater necessity. (B-C) Scatter plots show the parameters of thermodynamic models derived from CRISPRa (activation) and Enhancer-i (interference) for *MMP17* (B) and *CISH* (C). Parameters are shown as mean ± 95% confidence intervals. (D) Clustered correlation matrix showing Pearson correlations between potential predictors and both dCas9-VP16(10x) activation and dCas9-p300(core) activation. (E) Analysis of relative importance of predictors for activation using the LMG method (see methods). Importance is normalized to sum to 1.

To determine what genomic traits are predictive of enhancer strength, we looked at a set of 8 possible predictors of activation by both dCas9-p300(core) and dCas9-VP16(10x). The strongest correlated variables to dCas9-VP16(10x) and dCas9-p300(core) activation included the number of transcription factors (TFs) present at each site, the base level of RNAPII ChIP-seq signal and the base level of H3K27ac ChIP-seq signal (Figures 5D, S5E-H). Using a multiple regression-based method, the relative importance of each variable was calculated (see methods) (Groemping, 2006). This revealed that the best predictor of dCas9-VP16(10x) activation was the number of TFs present at the site, while the best predictor for dCas9-p300(core) mediated activation was the base amount of RNAPII present at each site (Figure 5E). The number of TFs bound to each site was determined by totaling the number of ChIP-seq peaks that overlap ERBS and adjacent regions using publicly available ChIP-seq data (ENCODE Project Consortium, 2012) for 18 different TFs in Ishikawa cells (Tables S2 and S3). While no TF was bound solely to strongly activated sites, we found certain TFs, such as TCF12 and ZBTB7A, were bound more often to enhancers that exhibited strong activation when targeted with synthetic activators (Table S2). Overall, these data suggest that dCas9-VP16(10x) and dCas9-p300(core) may have different requirements for target gene activation.

While the necessity and sufficiency of individual ERBS was consistent, combinations of ERBS behaved differently when comparing necessity and sufficiency. We previously reported synergy between ERBS when testing necessity. To quantitatively compare ERBS interactions between CRISPRa and Enhancer-i, we thermodynamically modelled the Enhancer-i data using the same model as activation with the difference being a site was defined as “active” if it was not blocked (i.e. untargeted by SID-dCas9-KRAB). In contrast to the models based on CRISPRa, we observed the expected cooperative interactions between pairs of ERBS: MMP17-1 and MMP17-2; CISH-1 and CISH-2; and FHL2-1 and FHL2-2 (Figure S5A-C). When comparing parameters between the Enhancer-i and CRISPRa derived models, it is clear that most parameter estimates are consistent, except ERBS interaction terms between activated enhancers (Figures 5B,C and S5D). These results are consistent with synergy in gene regulation occurring between ERBS when ER is binding to the enhancers, while ERBS that are instead bound by synthetic activators act independently on gene expression. Therefore, synergy likely occurs in the recruitment of ER and potentially cofactors to an ERBS, while targeting an activator directly to an ERBS does not require synergy with neighboring ERBS.

## Discussion

CRISPR-based gene activation is a unique platform for interrogating the sufficiency of gene expression enhancers within the native genomic context. In this study, we applied variations of the approach to distal regulatory elements normally bound by ER, using cells that were not exposed to estrogens and therefore had negligible ER activity. Targeting dCas9, fused to either the core of p300 or 10 copies of VP16, to 12 ERBS surrounding 4 genes simultaneously activated transcription to levels which mimicked an E2 transcriptional response. We also observed H3K27ac deposition at all sites targeted, in a pattern similar to that caused by E2 treatment. Regions adjacent to ERBS do not activate gene expression when targeted, suggesting that certain loci have the potential to impact gene expression when locally activated and others do not

In agreement with the idea of inherent activation potential of an enhancer, we also found that the sufficiency of an ERBS, as measured by dCas9-VP16(10x) gene activation from the site, is correlated with the necessity of the site, as measured by SID(4x)-dCas9-KRAB interference (Carleton et al., 2017). This observation supports the notion that inducible enhancers have an intrinsic ability to activate gene expression to a certain magnitude. Analysis of potential genomic factors which may determine activation potential revealed that the best predictor of dCas9-VP16(10x) activation is the amount of other transcription factors present at the site. A battery of other transcription factors may help stabilize dCas9-VP16(10x) or may perform orthogonal roles in enhancer maturation. This finding indicates that the underlying DNA sequence may be important for regulation by synthetic activators as they may be working together with other TFs. For dCas9-p300(core), the most important predictor is the base amount of RNAPII already present at the site, which could point to an important difference between the synthetic activators in their ability to recruit RNAPII, as VP16 has been shown to recruit basal transcription factors (Hirai et al., 2010). The unique predictors from this analysis suggests that different modes of activation are used by dCas9-VP16(10x) and dCas9-p300(core), which warrants further study.

We tested two dCas9 fusions and found that dCas9-VP16(10x) activates genes to a higher level than dCas9-p300(core). The fusions caused similar levels of H3K27ac to be deposited at targeted loci, suggesting that histone acetylation is not the only event that impacts transcription and that VP16 is likely contributing to gene activation in other ways. This could be because VP16 recruits a host of cofactors, including basal transcription factors and mediator, in addition to histone acetyl transferases (HATs) (Hirai et al., 2010). These interactions allow VP16 to more directly assemble the transcriptional machinery while p300-induced acetylation may be limited by other methods of transcriptional control, such as protein recruitment. The superior performance of dCas9-VP16(10x) may also be specific to the ERBS that we targeted, as dCas9-p300(core) has been shown to be more effective than VP16(4x) at other loci (Hilton et al., 2015), or it could be explained by the extra copies of VP16 in the 10x fusion.

While the necessity and sufficiency of individual ERBS was well correlated, the manner in which they combine to regulate gene expression was very different when tested with Enhancer-i and CRISPRa. In necessity experiments, pairs of ERBS showed synergistic behavior. For example, *CISH* was not estrogen responsive unless both CISH-1 and CISH-2 were active. In the sufficiency experiments, ERBS combined in a mostly independent/additive fashion, although some combinations appear sub-additive and may approach a saturating level of activation for the gene. One explanation for the independence between enhancers when bound by synthetic activators is that the synthetic activators are so strong that they override more subtle regulatory events that require synergy. However, we do not believe this to be the case, since an E2 induction leads to higher expression levels than either activator, especially dCas9-p300(core). The contrast in how ERBS combine to regulate transcription in the two experimental approaches suggests that the synergy between ERBS likely occurs in *cis*, where the recruitment of ER and its cofactors, such as HATs (Hanstein et al., 1996; Shang et al., 2000), is synergistic between ERBS. However, if the cofactors are directly recruited to the ERBS, as is the case with CRISPRa, then synergy no longer occurs. In this model, synergy occurs when ERBS influence one another prior to activation. Then, once ERBS are activated through the binding of transcription factors and cofactors, ERBS communicate with the target gene independently. There are important caveats to consider when using these synthetic activators including the possible recruitment of cofactors that do not normally bind to a particular enhancer or potential interference with TF binding by dCas9. However, we believe that the comparison between enhancer activation and enhancer interference has shed light on the consistent importance of individual sites as well as a key difference in how enhancers work together when bound by different transcriptional activators.

## Methods

### dCas9 Construct Generation

The Addgene 48227 plasmid (a gift from Rudolf Jaenisch) (Cheng et al, 2013) containing dCas9-VP16(10x) with a P2A linker and neomycin resistance gene was used for dCas9-VP16(10x) as well as the starting point for our dCas9-p300(core) construct. The p300(core) insert was obtained via PCR from the Addgene 61357 (gift from Charles Gersbach) (Hilton et al, 2015) plasmid using primers (Table S6), which also added AscI and ClaI restriction enzyme sites. Both the p300(core) PCR product and the dCas9-VP160 plasmid were digested by AscI and ClaI, removing the C-terminal VP16(10x) from the plasmid, and subsequently ligated together. Constructs were verified via Sanger sequencing (Table S6) (Genewiz).

### Guide RNA design

Guide RNAs (gRNAs) were designed and cloned as previously described (Carleton et al., 2017). 4 gRNA oligos were designed for each target region and pooled before Gibson cloning. gRNA plasmids were then pooled equally by site into three pools, such that each pool contained gRNAs targeted to one ERBS near each of the four genes studied as listed below. An adjacent pool was also created containing all gRNAs targeted to non-ERBS regions. Pools were created using equal mixtures of gRNA plasmids by mass.

**Pool 1:** MMP17-2, CISH-1, FHL2-2, HES2-3

**Pool 2:** MMP17-1, CISH-2, FHL2-3, HES2-1

**Pool 3:** MMP17-3, CISH-3, FHL2-1, HES2-2

**Adjacent Pool:** MMP17-A, MMP17-B, CISH-A, CISH-B, FHL2-A, FHL2-B

The sequences of the gRNAs targeting the ERBS surrounding *MMP17*, *CISH*, *FHL2*, *HES2*, adjacent regions, and promoters can be found in Table S4. Previously described gRNAs targeting the *IL1RN* promoter (Perez-Pinera et al., 2013) were used as controls.

### Cell culture and transfection

A human endometrial adenocarcinoma cell line, Ishikawa (Sigma), was used for ChIP-seq and gene expression experiments. Ishikawa cells were cultured in RPMI (Gibco) supplemented with 10% fetal bovine serum (Gibco) and 1% penicillin-streptomycin (Gibco) and incubated at 37°C with 5% CO_2_. Cells were transferred to hormone depleted media (phenol red-free RPMI (Gibco) with 10% charcoal-dextran stripped fetal bovine serum (Sigma)), at least 5 days before transfection by gRNA and dCas9 fusion plasmids. Ishikawa deletion lines were previously created and verified. All deletions are homozygous deletions selected through single cell cloning. Deletions were cultured in the same conditions as parental Ishikawa cells, again being transferred to hormone depleted media at least 5 days before transfection.

Cells were transfected using the Fugene HD reagent (Promega) according to the manufacturer’s protocol for unlisted cells. dCas9 fusions (dCas9-VP16(10x) or dCas9-p300(core)) plasmids were transfected at a mass ratio of 3:2 to pooled gRNA plasmids. Mass ratio of dCas9 fusions to tomato reporter plasmid was 6:1. Plasmid solutions were prepared in OptiMEM.

### ChIP-seq

Cells were grown in hormone depleted media for 5-7 days then plated in 15cm dishes at 8.5 million cells per dish. Cells were transfected 1 day after plating as described above with 42.75 μg total plasmid. Approximately 40 hours after transfection, media was changed to fresh hormone depleted media containing 1 μg/mL puromycin and 300 μg/mL G418 to select for cells transfected with both plasmids. Chromatin was harvested 72 hours post-transfection. ChIP was performed as previously described (Reddy et al., 2009). The antibodies used were HA (Biolegend 16B12), H3K27ac (Active Motif pAb Cat #39133) and RNAPII (abcam ab5408 [4H8]). Libraries were sequenced on the Illumina HiSeq 2500 as single-end 50 base pair reads and aligned to hg19 using bowtie with parameters -m 1 ‒t ‒best -q -S -l 32 -e 80 -n 2 (Langmead et al., 2009). Duplicates were removed from BAM files using samtools rmdup with modifier flag ‒s for single end reads (Li et al., 2009). Counts were generated using bedtools coverage (Quinlan and Hall, 2010) and normalized for total read depth. For H3K27ac ChIP-seq and RNAPII ChIP-seq, counts were then normalized to the average read depth within all overlapping peaks for a given antibody. H3K27ac levels before normalization show the same activation trends, with different baseline levels (Figure S2G). For HA ChIP-seq, there were not any overlapping peaks that were not targeted, therefore counts were normalized to the mean of 5 control regions with background signal: *CTCF* promoter (chr16:67,594,830-67,596,830), *TBP* promoter (chr6:170,862,978-170,864,978), *SF3B4* promoter (chr1:149,898,675-149,900,675), and *TRIM28* promoter (chr19:59,054,414-59,056,414). Fold change in ChIP-seq signal was calculated as the ratio of a targeted region’s normalized counts vs the same region’s normalized counts in a non-targeted control. ChIP-seq data for H3K27ac (DMSO and following an 8-hour E2 induction) was previously published and accessible at GEO (GSE99906) (Carleton et al., 2017).

### Gene expression analysis

Prior to gene expression analysis using qPCR, Ishikawa cell lines were grown in hormone depleted media for 5-8 days before being plated in 24 well plates at 60,000-100,000 cells per well. Cells were transfected 1 day after plating with 550ng total DNA per well. Approximately 40 hours after transfection, media was changed to hormone depleted media containing 1 μg/mL puromycin and 300 μg/mL G418 to select for successfully transfected cells. E2 inductions were performed by adding 10 nM E2 to media 64 hours post-transfection. 72 hours post-transfection, cells were lysed using Buffer RLT Plus (Qiagen) with 1% beta-marcaptoethanol (Sigma). RNA was isolated using the ZR-96-well Quick-RNA kit (Zymo Research) and quantified using either RiboGreen (Life Technologies) with an EnVision plate reader (PerkinElmer) or with a Qubit 2.0 (Life Technologies).

qPCR was conducted using the Power SYBR Green RNA-to-CT 1-Step Kit (Life Technologies). 50 ng of starting material was mixed into a 20 μL reaction volume. A CFX Connect light cycler (BioRad) was used to perform a 30-minute cDNA synthesis at 48°C followed by a 10-minute enzyme activation at 95°C and 40 cycles of 15 seconds at 95°C and 1 minute at 60°C. qPCR primers were added at a final concentration of 0.5 nM. Primer sequences are listed in Table S5. Primer specificity was confirmed using melt-curve analysis. Biorad CFX Manager 3.1 was used to calculate cycle threshold values using baseline subtracted curve fit and an auto-calculated single threshold. Final results were calculated using the ΔΔCt method with *CTCF* expression as the control. At least 2 replicates were analyzed per 24-well plate and at least 2 24-well plates were analyzed per experiment.

### Relative importance of predictors

Expression data and predictors were scaled to z-scores across all sites in R using the scale function with default parameters before analyzing Pearson correlation. To calculate relative importance, a regression for gene expression changes by dCas9-VP16(10x) and dCas9-p300(core) were fit based on 8 possible predictors using the lm function in R. Then, using the relaimpo package (Groemping, 2006), the relative importance of each predictor was calculated using the method described by Lindeman, Merenda and Gold (LMG) (Lindeman et al., 1980). Importance was normalized to sum to 1 for comparison. Distances to the nearest enhancers were calculated using H3K27ac peaks which do not overlap the TSS, as a proxy, using bedtools closest (Quinlan and Hall, 2010).

### Thermodynamic modelling

We used a modified version of the statistical thermodynamic model implemented in Gertz et al. (Gertz et al., 2009), originating from the Shea-Ackers formalism (Shea and Ackers, 1985).

For a system state s, the relative energy of that state is:

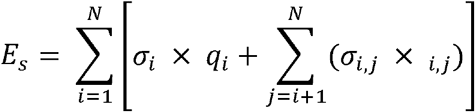

where N is the number of ERBS plus 1 for the promoter, M is the number of possible synthetic activator-synthetic activator interactions (or ER-ER interactions for the necessity data) and synthetic activator-RNAPII interactions, σ is a binary variable that denotes whether an ERBS or promoter is bound (0 for unbound and 1 for bound), q values represent protein:DNA interactions, and ω values represent protein-protein interactions. While relative q values are often fit using position weight matrices, in this case guide RNAs determine binding and we therefore leave the q values as free parameters.

We then calculate a thermodynamic weight for each state s as:

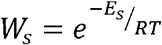

where R is the gas constant and T is temperature set at 37°C.

We define an experimental state, conditional on which enhancers are being targeted, which can be thought of as the union of possible states. For example, if sites 1 and 2 are being targeted, possible system states include: site 1 is bound, site 2 is bound, both are bound or none are bound; however, states with site 3 bound are considered highly unlikely and not considered.

Therefore, the probability of RNAPII being in a certain experimental state e is given by the partition

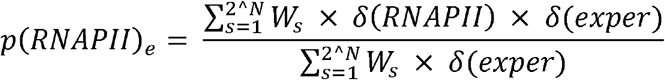

where δ(RNAPII) is a delta function which is equal to 1 when RNAPII is bound in a given system state and 0 otherwise. δ(exper) is a delta function corresponding to whether a system state is possible given which regulatory regions are being targeted in the experimental state. 2^N^ is the number of possible experimental states given N regulatory regions.

Gene expression was assumed to be correlated to the probability of RNAPII being bound to the promoter. When fitting our parameters, we maximized the value of the following correlation:

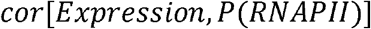

where cor is the Pearson correlation across all experimental states tested.

Random starts were chosen as a set of 10 random numbers between the limits of −5 and 5 using the runif function in R. In order to control for potential bias resulting from the optimization algorithm used to fit the parameters, parameters were fit using both the L-BFGS-B algorithm (Byrd et al., 1995), which is a gradient descent-based method and the Hooke and Jeeves Pattern Search Optimization method (Hooke and Jeeves, 1961) which does not rely on gradient descent. An interface to these algorithms was implemented in R by John C Nash in the optimr package (Nash, 2016). Using both algorithms, 1000 iterations were run, resulting in 2000 parameter fits. We then selected parameter sets which correlated with gene expression data reasonably well (within 0.1 of an optimal correlation coefficient) for downstream analysis and plotting.

Inhibition by SID(4x)-dCas9-KRAB was modeled in the same way with the exception that sites not targeted were defined as “active.” We therefore assume that if a site is targeted with SID(4x)-dCas9-KRAB it is completely inactive. The model is again fit on correlation between RNAPII occupancy and previously measured qPCR data (Carleton et al., 2017).

## Data Availability

The ChIP-seq data is available at the Gene Expression Omnibus (GEO) under accession GSE133300.

## Author Contributions

Conceptualization, M.G. and J.G.; Methodology, M.G., J.C., and J.G.; Investigation, M.G., J.C., A.R., and K.B.; Writing – Original Draft, M.G. and J.G.; Writing – Review & Editing, all authors; Funding Acquisition, J.G.

## Acknowledgements

This work was supported by NIH/NHGRI R01 HG008974 to J.G., and the Huntsman Cancer Institute. Research reported in this publication utilized the High-Throughput Genomics Shared Resource at the University of Utah and was supported by NIH/NCI award P30 CA042014. We thank Jeffery Vahrenkamp for analysis advice and we thank K-T Varley as well as Gertz and Varley lab members for their valuable comments on the research and the manuscript.

---

**Supplementary Figure 1.**
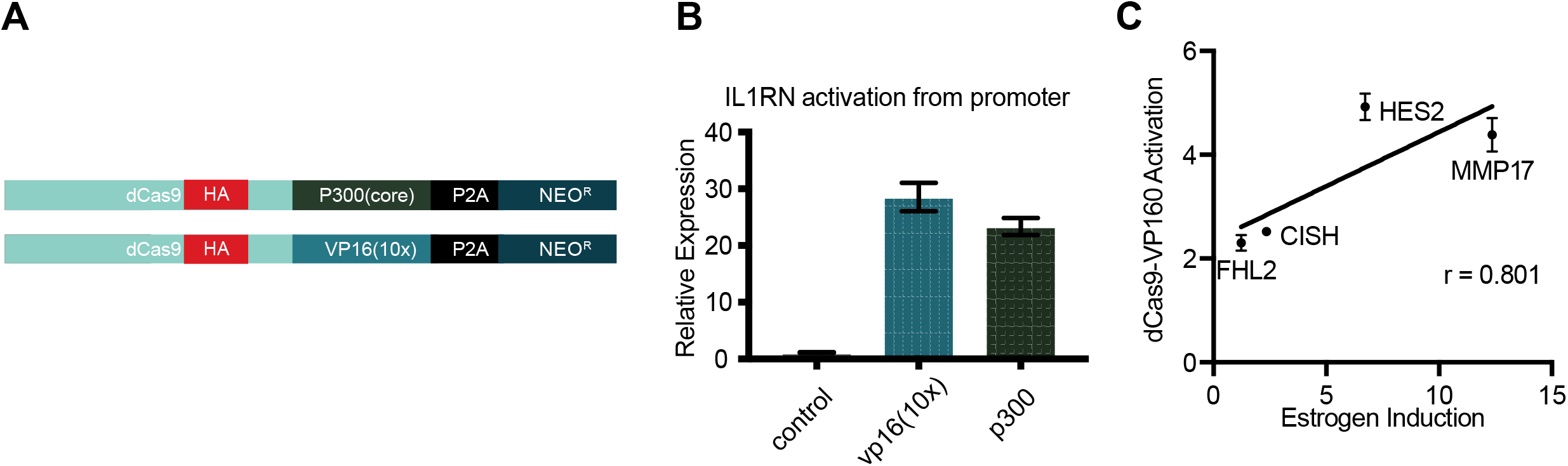
dCas9-activator constructs can activate gene expression (related to Figure 1). **(A)** A schematic shows dCas9-p300(core) and dcas9-vp16(10x) constructs used in this study. **(B)** *IL1RN* activation is induced by targeting either dCas9-p300(core) or dCas9-vp16(10x) to the *IL1RN* promoter. Error bars represent SEM. **(C)** Activation from simultaneously targeting all 3 ERBS near target genes induces activation to a level correlated with an 8-hour estrogen induction.

**Supplementary Figure 2.**
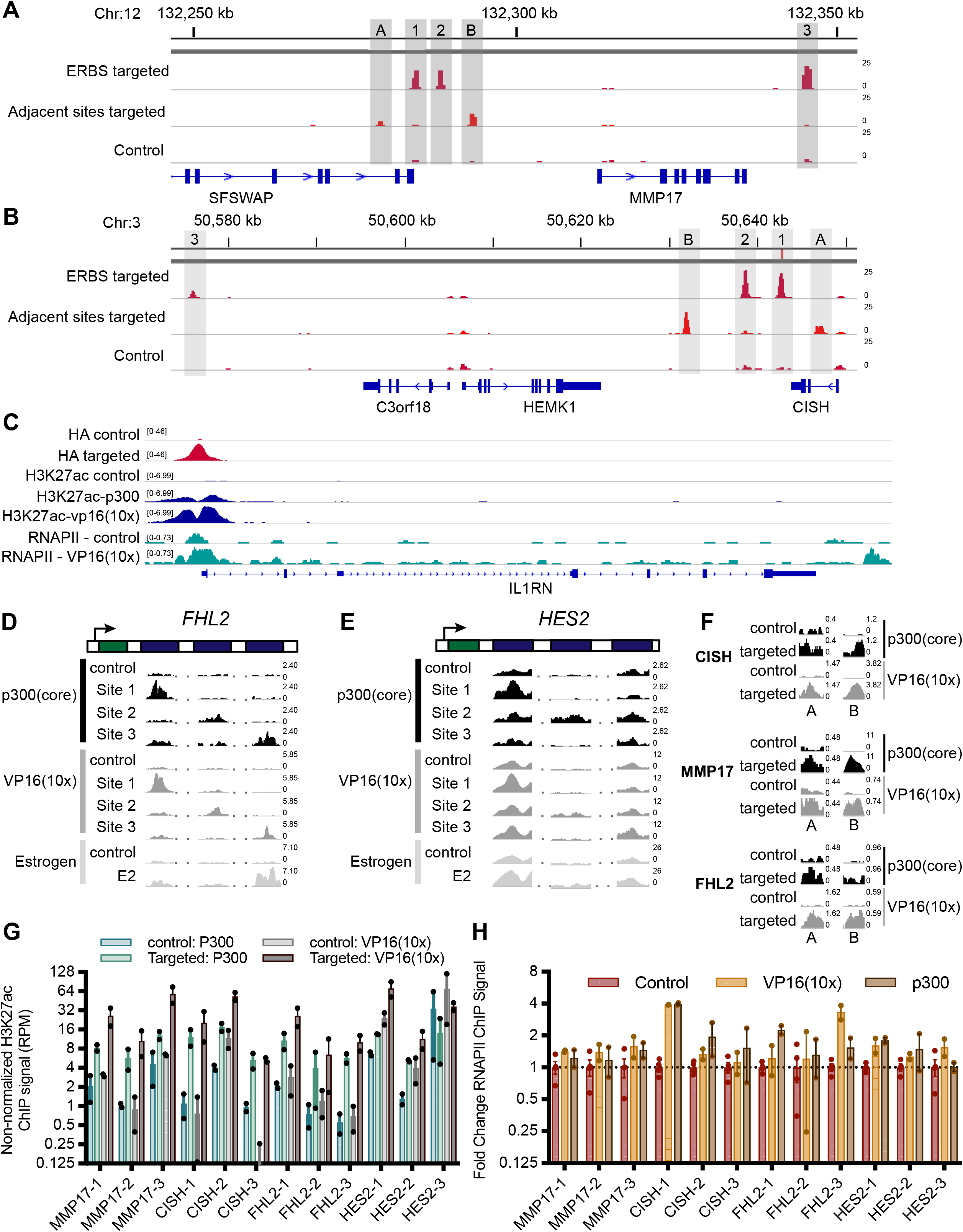
Targeting dCas9-VP16(10x) to ERBS and adjacent regions is specific and induces H3K27ac at the intended loci (related to Figure 2). **(A-B)** HA ChIP-seq browser tracks show specific targeting of ERBS (lane 1) and adjacent regions (lane 2) compared to a control with the *IL1RN* promoter targeted (lane 3) for the *CISH* **(A)** and *MMP17* **(B)** locus. **(C)** ChIP-seq browser tracks show targeting of dCas9 constructs to the *IL1RN* promoter (HA – lanes 1 and 2), H3K27ac induced by dCas9-p300(core) (lanes 3 and 4) or dCas9-vp16(10x) (lanes 3 and 5), and RNAPII induced by dCas9-vp16(10x) (lanes 6 and 7). **(D-E)** H3K27ac ChIP-seq browser tracks for sites targeted by dCas9-p300 and dCas9-vp16(10x) show similar histone acetylation as an 8-hour estrogen treatment for *FHL2* **(D)** and *HES2* **(E)**. **(F)** Targeting dCas9-p300(core) or dCas9-VP16(10x) to regions adjacent to ERBS induces H3K27ac. **(G)** H3K27ac levels before normalization (shown as reads per million) for ERBS before and after targeting by dCas9-p300(core) (blue/green) or dCas9-VP16(10x) (gray) **(H)** Limited fold changes in RNAPII levels were observed when targeting individual ERBS with dCas9-VP16(10x) or dCas9-p300(core).

**Supplementary Figure 3.**
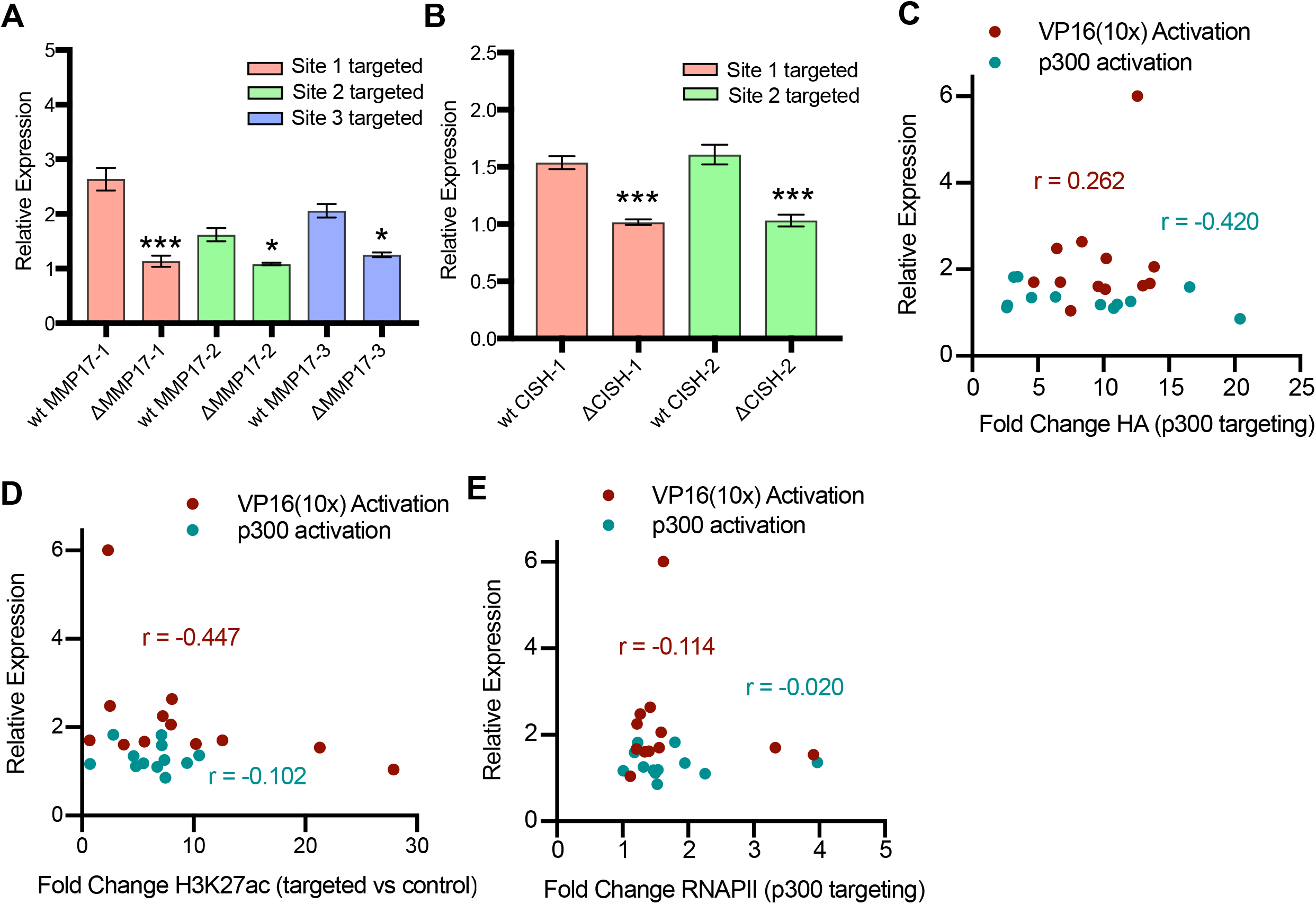
Activation from individual ERBS is specific as well as independent of targeting efficiency and acetylation induced by dCas9-vp16(10x) (related to Figure 2, Figure 3 and Figure 5). **(A-B)** dCas9-VP16(10x) was targeted to ERBS in clonal cell lines in which individual ERBS have been deleted. When targeting the deleted site at ERBS surrounding *MMP17* **(A)** and *CISH* **(B)**, we do not see activation, confirming specificity of activation. P-values are calculated using a holm adjusted t-test (***: p-value < 0.001,**: p-value < 0.01, *: p-value < 0.05). **(C-E)** Correlation plots showing relationship between activation by dCas9-p300(core) **(teal, C-E)** or dCas9-VP16(10x) **(red, C-E)** and fold change ChIP-seq signal of HA (dCas9) **(C)** H3K27ac **(D)** and RNAPII **(E)**. r is Pearson’s correlation coefficient.

**Supplementary Figure 4.**
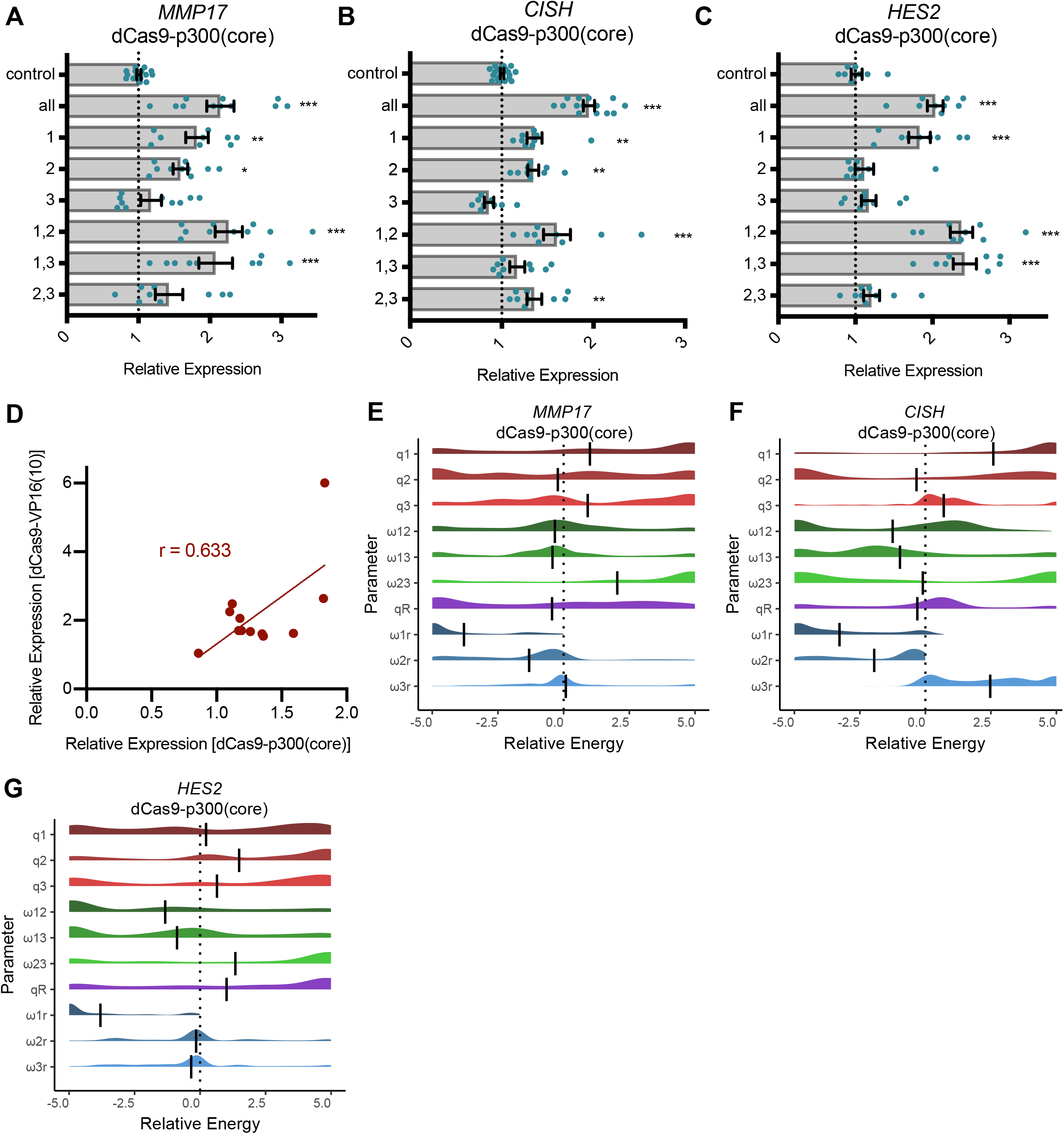
Thermodynamic modelling of dCas9-p300 shows independence between sites (related to Figure 3 and Figure 4). **(A-C)** Relative fold change expression levels from targeting combinations of ERBS with dCas9-p300(core) as measured by qPCR for *MMP17* **(A)** *CISH* **(B)** and *HES2* **(C)**. **(D)** Correlation between activation by dCas9-VP16(10x) and dCas9-p300(core) at individual ERBS. **(E-G)** Parameters of a thermodynamic model were fit to relative expression data from combinatorial activation of ERBS with dCas9-p300. **(E)** Interactions between sites at *MMP17* remain mostly neutral. *CISH* **(F)** and *HES2* **(G)** interaction parameters appear bimodal.

**Supplementary Figure 5.**
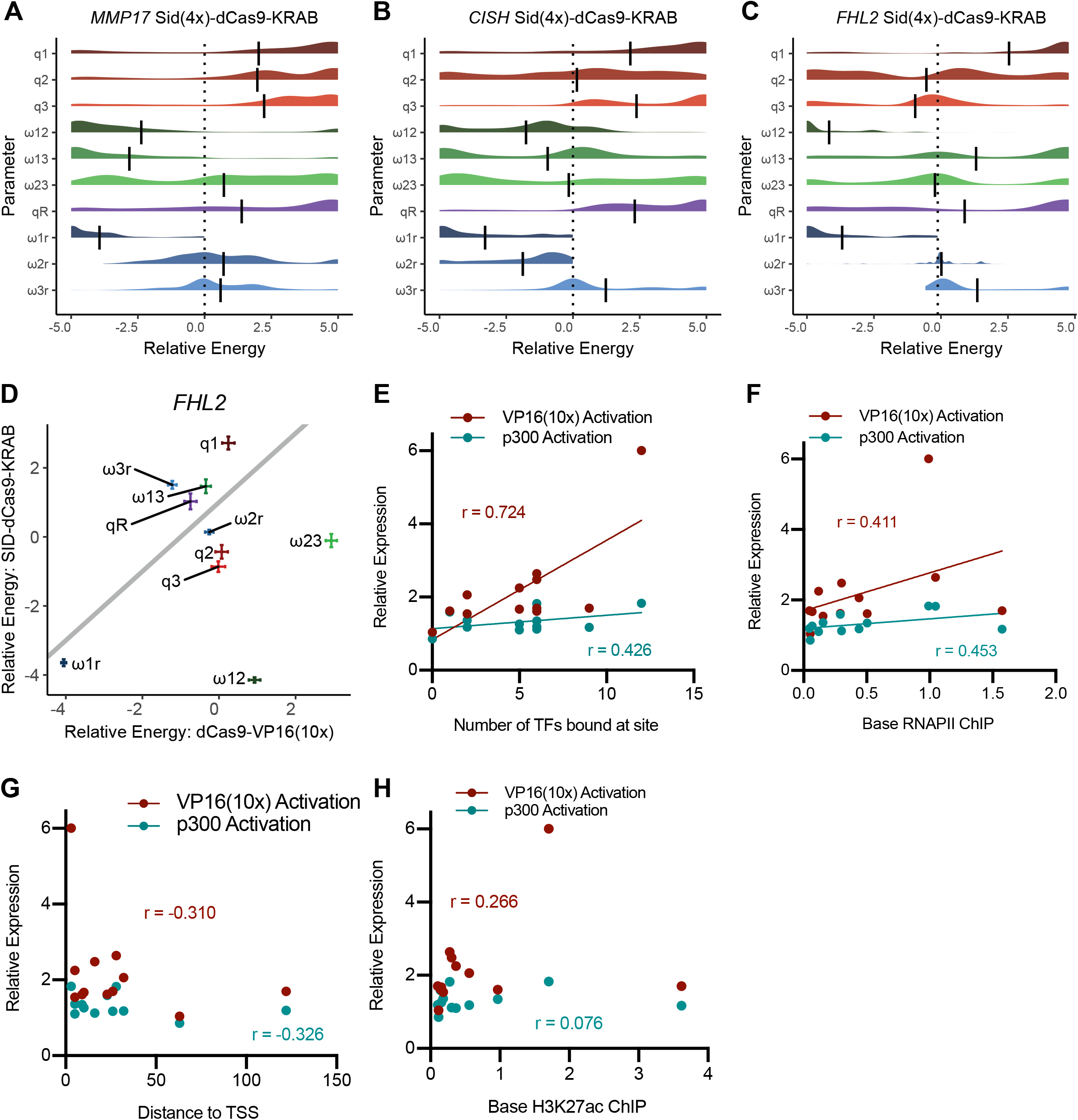
Thermodynamic modelling of Enhancer-I shows cooperativity between sites and correlation of predictors with gene activation (related to Figure 4 and Figure 5) **(A-D)** Parameters were fit to relative expression data from combinatorial inhibition of ERBS with SID(4x)-dCas9-KRAB (Carleton et al., 2017). We see favorable (more negative) interactions between ERBS 1 and 2 (parameter ⍰_1,2_) for *MMP17* **(A)**, *CISH* **(B)** and *FHL2* **(C)**. Vertical bars represent the mean. **(D)** Comparison of modelled parameters from activation to inhibition showing mean fitted parameters in the two studies. We see that the cooperativity of FHL2-1 and FHL2-2 is specific to inhibition, while the remaining parameters are relatively similar. Error bars represent 95% confidence intervals around the mean. **(E-H)** Correlation plots showing relationship between synthetic activation dCas9-p300(core) (teal) or dCas9-VP16(10x) (red) and possible genomic predictors of activation (x-axis).

